# Impact of nanoscale hindrances on the relationship between lipid packing and diffusion in model membranes

**DOI:** 10.1101/2020.01.16.907816

**Authors:** Daniel Beckers, Dunja Urbancic, Erdinc Sezgin

## Abstract

Membrane models have allowed for precise study of the plasma membrane’s biophysical properties, helping to unravel both structural and dynamic motifs within cell biology. Free standing and supported bilayer systems are popular models to reconstitute the membrane related processes. Although it is well-known that each have their advantages and limitations, comprehensive comparison of their biophysical properties is still lacking. Here, we compare the diffusion and lipid packing in giant unilamellar vesicles, planar and spherical supported membranes and cell-derived giant plasma membrane vesicles. We apply florescence correlation spectroscopy, spectral imaging and super-resolution STED-FCS to study the diffusivity, lipid packing and nanoscale architecture of these membrane systems, respectively. Our data show that lipid packing and diffusivity is tightly correlated in free-standing bilayers. However, nanoscale interactions in the supported bilayers cause deviation from this correlation. This data is essential to develop accurate theoretical models of the plasma membrane and will serve as a guideline for suitable model selection in future studies to reconstitute biological processes.

## Introduction

The cascades for signal transduction usually begin at the cell surface, and for this reason the plasma membrane can be considered as the main hub for cellular signalling^1^. However, drawing conclusions about membrane behaviour and architecture proves challenging, not least because poorly understood or still unknown processes influence its dynamics^2, 3^. Our current knowledge shows that plasma membrane is a vastly complex and intricate system^4^. Therefore, to truly appreciate and understand the finesse behind membrane dynamics, a ‘bottom-up’ approach to discern different processes can prove useful^5^. Several systems address this, employing a basic skeleton of only the essential biological components of the plasma membrane but engineered to allow systematic incorporation of complexity^6, 7^. Such reductionist systems can not only mimic membranes, but also allow membrane associated events to be systematically broken down to reveal their key contributing species owing to their controllable compositional complexity^7^. Popular models include free-standing bilayers of synthetic lipids such as giant unilamellar vesicles (GUVs)^8^ or membrane blebs of live cells known as giant plasma membrane vesicles (GPMVs)^9, 10^. However, the development of solid substrates to support bilayers have also shown promise, with two prominent constructs being the planar substrate/supported lipid bilayers (SLBs)^11, 12^ and spherical bead supported lipid bilayers (BSLBs)^13, 14^ (also termed spherical supported lipid bilayers, SSLBs).

It is certain that membrane models will continue to aid our understanding of the dynamics that underlie cellular signalling. Though, it is worth to note that each model brings their advantages and limitations and caution should be employed while choosing appropriate model systems for given biological processes. It is, therefore, imperative to understand how each model influences bilayer behaviour, not only to best select appropriate models for future research, but to also avoid drawing misleading conclusions. This necessitates comprehensive comparison between models.

Here, we directly compare the biophysical properties of GUVs, SLBs, BSLBs and cell-derived GPMVs. We apply florescence correlation spectroscopy (FCS)^15^, spectral imaging^16^ and super-resolution STED spectroscopy^17^ to study the diffusivity, lipid packing and nanoscale architecture of these membrane systems, respectively. We observed slower diffusion for SLBs and BSLBs compared to GUVs as reported in the literature previously^18^. Whilst spectral analysis revealed no difference in lipid packing within these systems, STED combined with FCS showed nanoscale hindrances within SLBs and BSLBs that would explain their comparatively slower diffusion rates despite their similar lipid packing. Moreover, we showed that changes in lipid packing and diffusion in GUVs in response to compositional changes are predictable while support has significant influence on this relationship. This work highlights the necessity of carefully comparing membrane models to progress research in membrane biology.

## Materials and Methods

### Cell lines, lipids and dyes

Chinese hamster ovary (CHO) cells were cultured in DMEM/F12 media supplemented with 10% FBS and 1% L-glutamine. Cells were prepared two days prior to experiments. Lipids stocks were obtained from Avanti Polar Lipids. GUV, SLB, and BSLB bilayers were prepared to contain 1% DGS-Ni-NTA with 1 mg/ml lipid stocks of POPC (1-palmitoyl-2-oleoyl-sn-glycero-3-phosphocholine), POPC:Cholesterol (of varying concentrations) and DPPC (1,2-dipalmitoyl-sn-glycero-3-phosphocholine):Cholesterol (1:1), all in chloroform. Lipid stocks were stored under nitrogen at -20 °C. FCS and confocal imaging were performed with phosphatidylethanolamine (PE) labelled with Abberior Star Red (herein referred to as AbStR-PE) that is obtained by Abberior. Spectral imaging was performed with C-Laurdan obtained by 2pprobes.

### Generation of model membranes

GUVs were prepared by electroformation^19, 20^. With this approach unilamellar vesicles between 10-100 µm diameters in size are produced. Lipid stock was spread onto two parallel platinum wires attached to a custom-built Teflon-coated chamber and left briefly to evaporate solvent. Wires were passed under nitrogen gas before submersion in 300 mM sucrose. 10 Hz AC current was applied to wires for 1 hour to trigger vesicles swelling, followed by 2 Hz for 30 minutes.

GPMVs were prepared as described previously by Sezgin^10^. Briefly, CHO cells were grown to 60 % confluency, washed three times with GPMV buffer (10 mM HEPES, 2 mM CaCl_2_, 150 mM NaCl, pH 7.4) and then incubated at 37 ^o^C in GPMV buffer with PFA and DTT (10 mM HEPES, 2 mM CaCl_2_, 150 mM NaCl, 25 mM PFA and 2 mM DTT) for 1-2 hours to stimulate membrane blebbing. The resulting supernatant containing GPMVs was then extracted.

SLBs were prepared by spin-coating^21^. Briefly, glass coverslips of ϕ25 mm and #1.5 mm thickness were first cleaned in piranha-solution (sulfuric acid (95-98%): hydrogen peroxide (30%), 3:1) for 1 hour. Cleaned cover slips were then repeatedly washed and then stored in distilled water for no longer than 1 week. 25 µl of 1 mg/ml of lipid stock was pipetted onto the centre of a dried cover slip and immediately spun for 30 seconds at 3500 rpm. The coated coverslip was then placed into a metal chamber and rehydrated with 1 ml of SLB buffer to form a bilayer (150 mM NaCl, 10 mM HEPES, pH 7.4).

BSLBs were prepared from spontaneous fusion of liposomes of lipid stock with 5 µm silica beads^14, 22^ obtained from Bangs Laboratories. Liposomes were prepared by tip sonication. Lipid stock was placed under nitrogen gas to evaporate solvent completely leaving a dry, thin lipid film. Tris Buffer Saline solution (50 mM Tris HCL, 150 mM NaCl, pH 8.0) (500 µl) was added to lipid residue as liposome buffer. Lipid solution was then transferred to ice and sonicated at 55 Amp for 15 minutes, with 10 second pulse periods separated by 10 second rest intervals. Silica beads were washed with 1 ml PBS before centrifuging for 30 seconds at 2000 rpm. Supernatant was removed with residual left to prevent beads from drying, and washing was repeated twice. Beads were mixed with liposomes (1:5) and then shaken for 20 minutes at 1200 rpm to form BSLBs. After centrifuging at 2000 rpm for 30 seconds, BSLBs were then washed twice with PBS, with ∼500 µl solution reserved from the final wash.

### Confocal Imaging

Membranes were imaged with Zeiss LSM 780 or 880 microscopes. First, GUVs, BSLBs, and GPMVs were incubated with 0.1 µg/ml AbStR-PE for 15 minutes. GUV, BSLB, and GPMV were transferred to Ibidi 8-well plastic chambers of #1.5 thickness. Wells were previously treated with 1 mg/ml BSA, left for 1 hour and then washed three times with PBS or GPMV buffer before transfer. GUVs and BSLBs were then suspended in PBS. GPMVs were left unsuspended for 1 hour to allow to settle for imaging. SLB lipid mixture was prepared with 0.1 µg/ml AbStR-PE and imaged as is. 633 nm laser was focused onto bilayers by 40X water immersion objective (NA 1.2) for excitation of AbStR-PE fluorophore.

### Confocal and STED Fluorescence correlation spectroscopy (FCS)

FCS was used to measure and compare the diffusion of AbStR-PE through GUV, SLB, and BSLB models prepared with POPC, POPC:Chol, and DPPC:Chol compositions, and GPMVs. Models were incubated with 0.05 µg/ml AbStR-PE as previously described. GUVs, BSLBs, GPMVs were measured in Ibidi glass chambers of #1.5 thickness prepared as before. SLBs were measured on ϕ25 mm and #1.5 mm thickness glass cover slides.

Confocal FCS was performed on Zeiss LSM 780 microscope with 40X NA 1.2 water immersion objective. Before measurement, focal spots were calibrated using a mixture of Alexa 488 and 647. FCS measurements were recorded with a 633 nm laser at 0.1% power (≈2 µW). Laser focusing was completed by finding axial positions of maximum fluorescence intensity at the bilayer. Correlation curves were obtained over five second periods with five repeats per area studied. Curves were then fitted with the freely available FoCuS-point software to extract diffusion coefficients^23^. All FCS data were fitted with to a 2-dimensional diffusion model that incorporated an initial triplet state that describes a fixed 5 µs relaxation period. STED-FCS was performed with Leica SP8 microscope using 100X NA 1.4 oil immersion objective. Laser focusing and data acquisition was performed as previously described^24^.

### Spectral imaging

Spectral imaging was used to measure and compare packing within bilayers. GUVs, BSLBs, SLBs and GPMVs were incubated with 1 µM C-Laurdan for 10 minutes. GUVs, BSLBs, and GPMVs were then transferred to Ibidi plastic bottom chambers prepared as previously described. Imaging was performed by Zeiss LSM 780 microscope equipped with a 32-channel GaAsP detector array and a polarizer that minimizes photoselection effect. Laser light at 405 nm was selected for C-Laurdan excitation and the λ-detection range was set between 415 and 691 nm. Images were analysed with the custom GP plugin of FIJI software as described previously^16^.

## Results

There is already substantial evidence of a support’s influence on bilayer diffusion^25-28^. To further assess the influence of support on other pivotal biophysical parameters and the relationships between them, we have selected four models that broadly represent the spectrum of designs extensively utilised. Our chosen models span from free-standing GUVs, to supported planar SLB and spherical BSLB constructs, to the more cellular inspired bilayer model of GPMVs (Fig. 1). GUVs are free-standing vesicles, and for this reason exhibit considerable polydispersity ranging in diameters of 10-100 µm within populations. SLBs and BSLBs are by contrast supported on substrate. Not only does the substrate confer mechanical stability, but in the case of BSLBs offers the attractive option of size-tuning. Whilst BSLBs retain a spherical construct, SLBs lack three-dimensionality and instead model the bilayer as an infinitely flat construct. In both supported models, the substrate influence on bilayer behaviour, particularly its effect on diffusion, has been consistently reported^18, 29^. Finally GPMVs, like GUVs, are free-standing but derived from live cells^10^. As a result, their membrane composition reflects native cell character but is removed from the influence of an actin cytoskeleton. We measured the diffusion and lipid packing in these four prominent bilayer models through the application of FCS, spectral imaging, and super-resolution STED spectroscopy. More importantly, we assessed the correlation between lipid packing and diffusion within different models as well as different compositions.

**Figure 1.**
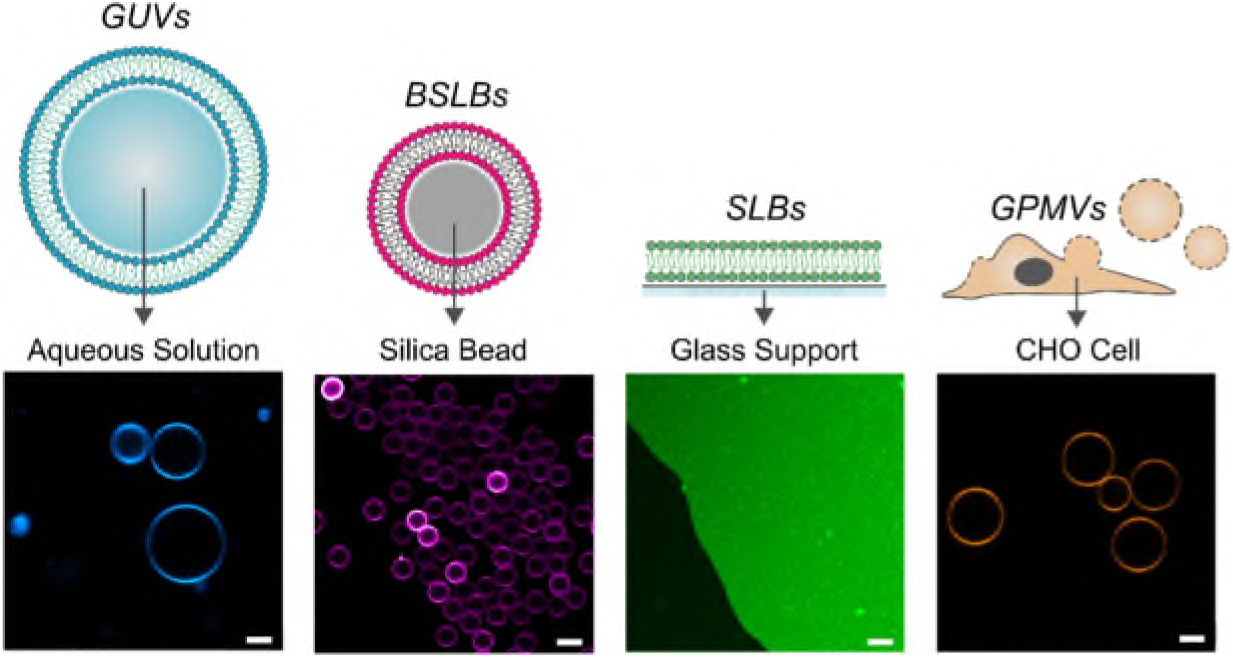
Illustrations and confocal images of membrane models. Illustrations highlight details in model designs. Confocal images were obtained at the equatorial plane of membranes labelled with AbStR-PE lipid fluorophore. All scale bars are 5 µm.

### GUV models confirm a relationship between lipid packing and diffusivity within the bilayer

We first considered the free-standing bilayer model of GUVs. GUVs are by definition free from a support influence and thus can reveal unbiased relationships between biophysical parameters of the bilayer. Here, we systematically altered the composition of GUVs in the form of glycerophospholipid species and cholesterol concentration with the intent of incrementally increasing bilayer ordering and seeing how this correlates with diffusion. C-Laurdan^30^ was incorporated into bilayers to report on lipid ordering^31^. Spectral imaging^16^ confirmed increased ordering as compositions progressed from comprising fully unsaturated lipid (DOPC, 18:1/18:1) to monounsaturated (POPC, 16:0/18:1), to saturated lipid (DPPC, 16:0/16:0) (Fig. 2A), and increased too alongside cholesterol concentration, as expected^32, 33^. Extracting generalised polarisation (GP) scores that quantify ordering within images revealed a monotonic increase in ordering as GUVs comprised with higher concentrations of cholesterol (Fig. 2B, violin plots). We then gauged for the influence of lipid composition on bilayer diffusivity. To this end, we employed point FCS to obtain the lateral diffusion coefficients of Abberior Star Red-labelled phosphatidylethanolamine (AbStR-PE) analogue incorporated within GUVs of the same compositions. Measurements for GUVs agree with earlier reports for other similarly structured fluorophores (Fig. 2B, box-and-whisker plots)^34^. Diffusion coefficients decreased with lipid saturation and also as a function of cholesterol concentration, agreeing with previous studies, and describes an approximately linear correlation^35^. Overlaying respective diffusion coefficients and GP scores revealed a negative correlation between the two measures and so suggests a distinct relationship between bilayer ordering and diffusivity in GUV models (Fig. 2B).

**Figure 2.**
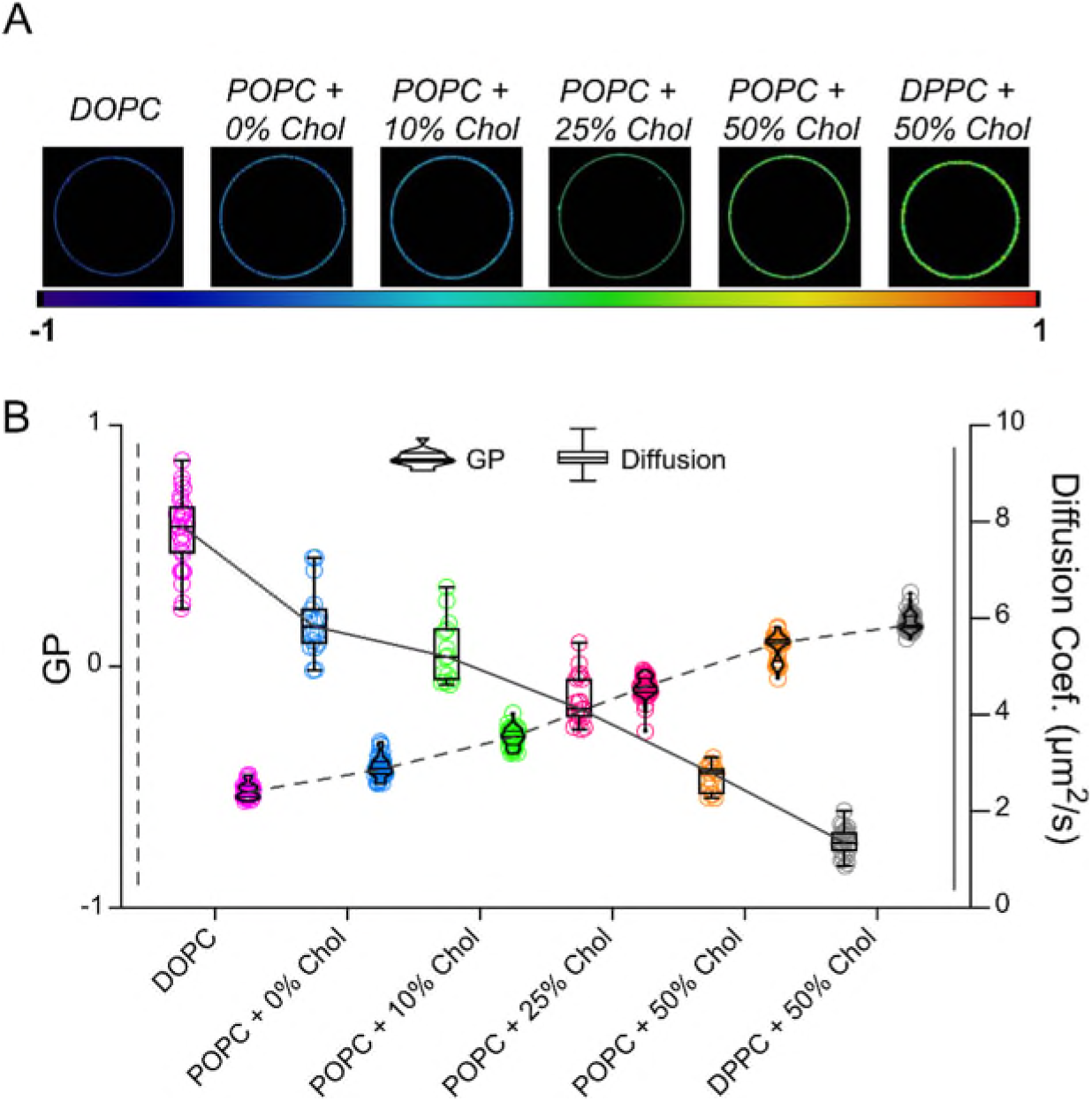
Relationship between lipid packing and diffusion in free-standing GUVs. (A) Spectral images of different GUV compositions. The colour code (below images) corresponds to packing and relates to GP values wherein higher values indicate tighter lipid packing. (B) C-Laurdan GP (violin plots) and AbStR-PE diffusion (box-and-whisker plots) measurements of GUVs across different compositions.

### Diffusion in GUVs differ from supported membrane models

Having observed a clear relationship between lipid ordering and mobility in GUVs, we sought to investigate whether this relationship extended to other models. As discussed, we selected models that together broadly capture current key design motifs; herein we investigated aforementioned GUVs alongside SLBs and BSLBs of supported planar and spherical designs respectively, as well as free-standing GPMVs that reflect the complex composition of native bilayers. First, we established diffusion profiles of all models in identical conditions for comparison. FCS was performed in models, for the interest of simplicity, composed of single-component POPC. Confocal images confirmed homogenous fluorescence signal throughout respective bilayers at the microscopic level (Fig. 1). FCS curves for all models fitted well to a one-component two-dimensional diffusion model (Fig. 3A). Expectedly, diffusion coefficients measured highest in GUVs which demonstrated an approximate three-fold increase in mean diffusion coefficient over SLBs and GPMVs, with an approximate five-fold increase over BSLBs. Our results generally agree with previous studies that separately demonstrated an increase in diffusion speeds in GUVs compared with in SLBs^18^ and GPMVs^36, 37^.

**Figure 3:**
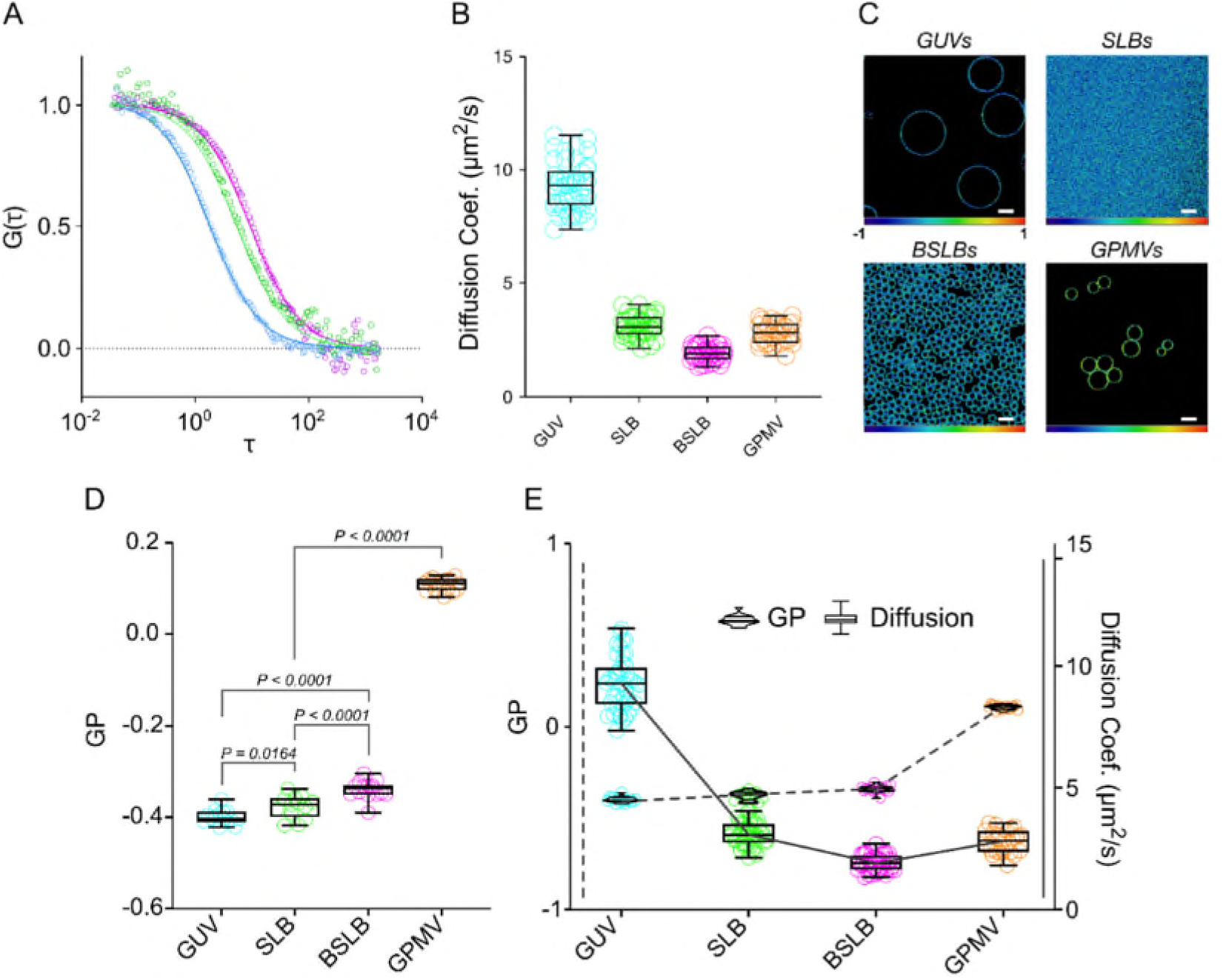
Diffusion and GP in membrane models. (A) Representative normalised point FCS curves of each membrane model composed of single-component POPC. (B) Diffusion coefficients of AbStR-PE through model bilayers. (C) GP maps of GUVs, SLBs, BSLBs, and GPMVs taken at their equatorial plane. Scale bars are 10µm. (D) GP values of membrane models. (E) Direct comparison of GP and diffusion coefficients of POPC bilayer models and GPMVs. Violin plots and box-and-whisker plots were assigned to GP and diffusion data, respectively.

### Lipid packing and diffusion do not correlate across different supported models

Having revealed the trends in diffusivity across models, we then set to investigate their lipid packing and assess for a correlation as observed previously in GUVs. Spectral imaging expectedly reported homogeneous packing within all four models, i.e., no notable microscopic inhomogeneity (Fig. 3C). Although statistically significant, there was an extremely small difference in packing between BSLBs, SLBs and GUVs (Fig. 3D). GPMVs, on the other hand, showed higher GP values compared to all model systems (Fig. 3D), presumably due to their complex lipid composition which includes saturated lipid components and cholesterol. When we overlaid GP data with diffusion coefficients, we observed no correlation between the two, suggesting the slow diffusion trends reported within supported models is not dictated by lipid packing (Fig. 3E).

### STED-FCS reveals nanoscale hindrances in the architecture of supported models

Until this point, diffusion measurements were performed on the diffraction-limited resolution scale which cannot report on nanoscale dynamics. In contrast, super-resolution STED combined with FCS (STED-FCS) offers resolution down to 20-40 nm (Fig. 4A) with which nanoscale dynamics can be addressed^17, 38^. In this regard, STED can glean assessment of bilayer dynamics in the context of surface architecture, an appreciated consideration in supported model design^27^. For instance, unwanted aggregates or surface defects could hinder diffusion of the fluorescent species as they travel through the observation spot and thus yield slow diffusion that is independent of the microscopic viscosity or lipid packing of the sample. Measuring the diffusion in a reduced focal volume with a STED laser allows us to distinguish free Brownian diffusion from hindered diffusion (Fig. 4A)^17, 38, 39^. A simple measurement yielding the ratio of confocal (*τ*_*C*_) and STED (*τ*_*S*_) transit time through the focal spot 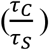 will reveal any differences (and hence unexpected hindrances) within the models. Nanoscale hindrances lead to a higher *τ*_*S*_ values, thus smaller 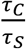 ratios.

**Figure 4:**
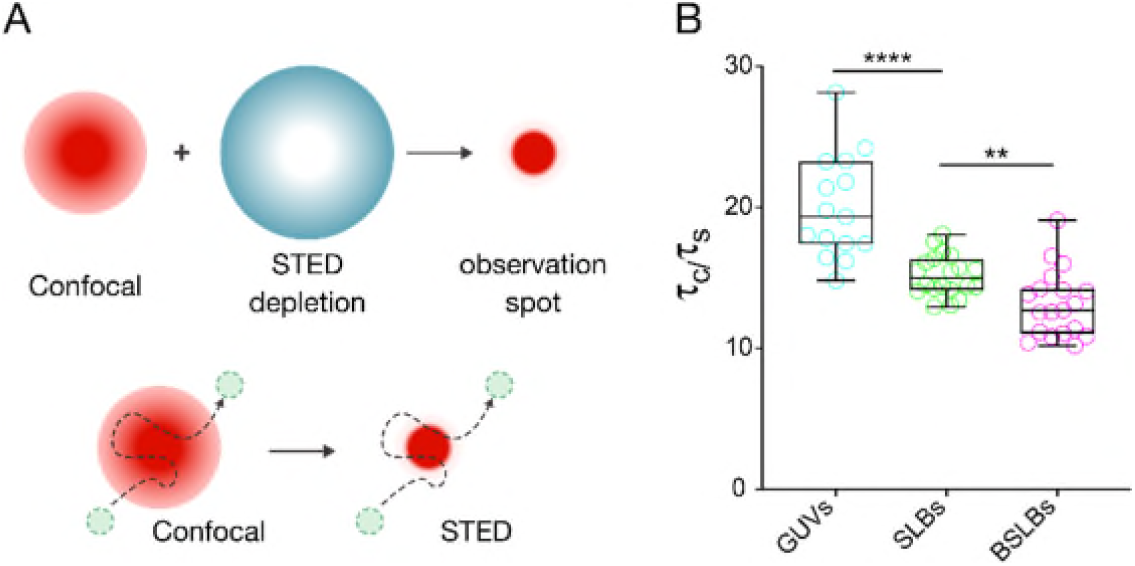
STED-FCS to reveal nanoscale hindrances in model systems. (A) Principle of STED-FCS. A super-resolved observation volume can be produced by a depletion beam (blue ring) designed with a zero intensity centre that effectively cancels surrounding emission signal from excited fluorophores. (B) Ratio of calculated confocal (τ_c_) and STED (τ_s_) transit times in model membranes composed of single-component POPC. A lower ratio of confocal:STED transit times suggests nanoscale hindrances.

We performed confocal FCS on model membranes composed of single-component POPC as before, immediately followed by STED-FCS. The ratio of confocal and STED transit times 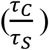 was calculated for all models (Fig. 5B). GUVs demonstrated a 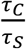 value of 19.7 ± 3.7 while SLBs and BSLBs showed 15.2 ± 1.4 and 13.1 ± 2.3, respectively. This suggests near free diffusion, i.e. minimal hindrance on lipid mobility in GUVs, but hindrances in SLBs and BSLBs at the nanoscale resulting in higher τ_S_ values.

**Figure 5.**
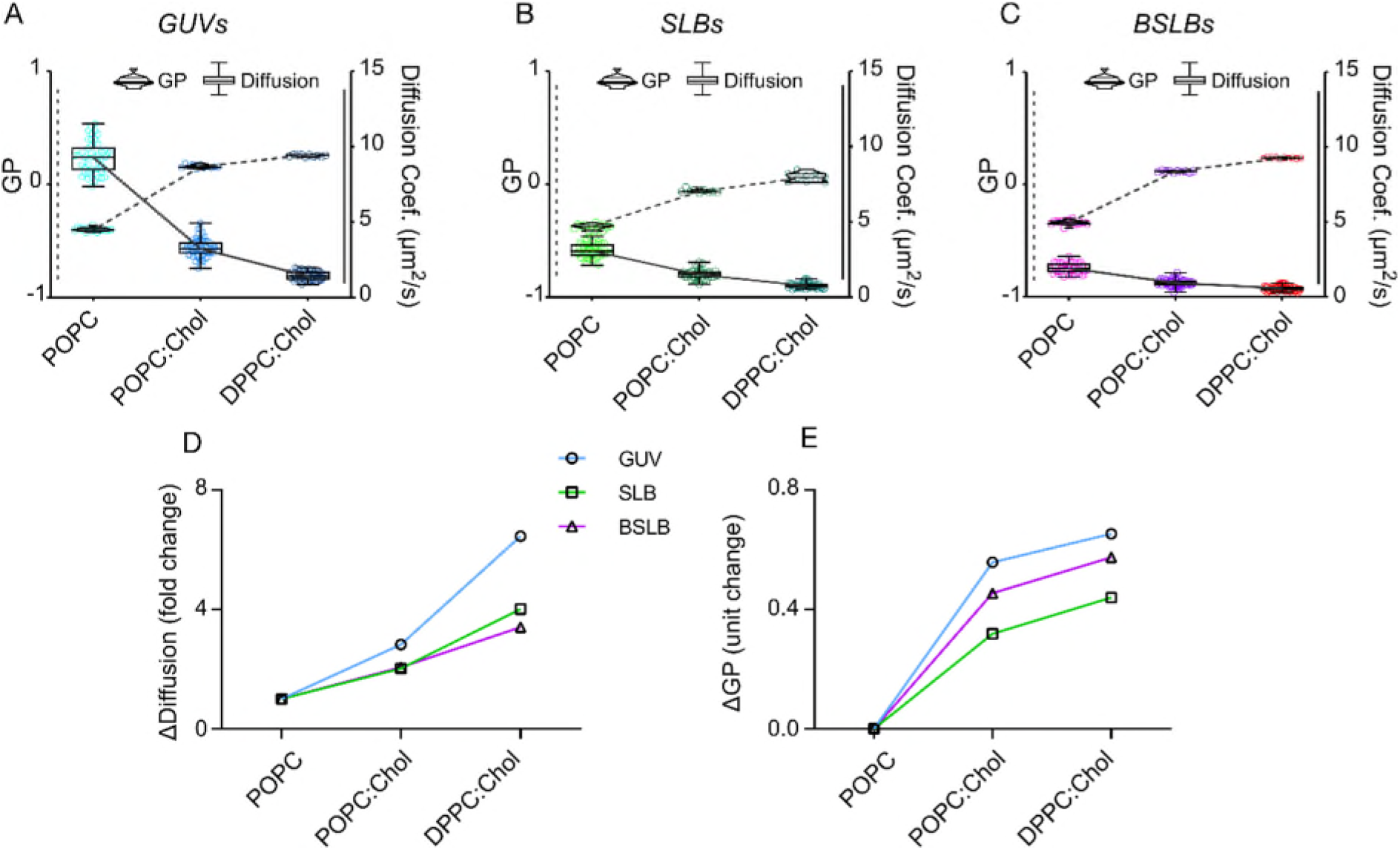
Relationship between lipid ordering and diffusion across lipid compositions. Direct comparison of GP (violin plots) and diffusion (box-and-whisker plots) within (A) GUV, (B) SLB, and (C) BSLB models of POPC, POPC:Chol and DPPC:Chol. Dotted lines indicate trends in GP whilst solid lines indicate trends in diffusion. (D) Calculated fold change in mean diffusion coefficient across compositions within models. (E) Calculated unit change in mean GP value across compositions within models.

### The relationship between lipid ordering and diffusion is less pronounced in supported models

We have shown that lipid packing is comparable in bilayers of our single component POPC models (Fig. 3D), but supported models exhibit altered diffusion profiles and hint towards a ‘broken’ relationship between ordering and diffusion in such models. We sought to demonstrate how this discrepancy is affected in more ordered membrane systems by incrementally increasing ordering in GUVs, SLBs, and BSLBs. We then measured GP and diffusion and overlaid them. GP analysis of spectral images confirmed an increase in ordering as bilayer compositions changed from POPC to POPC:Chol (1:1), and DPPC:Chol (1:1) (Fig. 5A-C). Confocal FCS measurements for each model also showed an expected trend for all models; slower diffusion for more saturated membranes. (Fig. 5A-C).

To quantitatively assess how these two parameters are correlated and to have a more comprehensive picture of the changes in diffusion and GP at different compositions, we calculated the “fold change” in diffusion (mean diffusion coefficients in POPC divided by mean diffusion coefficient in POPC:Chol or DPPC:Chol). Similarly, we calculated unit change in GP (mean GP in POPC:Chol or DPPC:Chol minus mean GP in POPC) for all model systems (Fig. 5D, E). As expected, more ordered membrane systems yielded slower diffusion and this trend was maintained in all systems (Fig. 5D, E), but importantly to different extents. In GUVs, the differences in diffusion as well as GP (between ordered and disordered membrane systems) is the highest. This suggests that free-standing GUV system is very sensitive to compositional changes. However, SLBs and BSLBs did not react as well to the changes in saturation. For instance, ΔDiffusion value for DPPC:Chol (i.e., mean diffusion coefficient of fluorescent lipid in POPC divided by mean diffusion coefficient of fluorescent lipid in DPPC:Chol) was ≈6.5 for GUVs while it was ≈4 and ≈3.4 for SLBs and BSLBs, respectively. In other words, diffusion in DPPC:Chol GUVs is 6.5 times slower compared to POPC GUVs; however, diffusion in DPPC:Chol SLBs is only 4 times slower compared to POPC SLBs (Fig. 5D).

Similarly, ΔGP for DPPC:Chol (i.e., mean GP in DPPC:Chol minus mean GP in POPC) is ≈0.65 units for GUVs, and ≈0.44 and ≈0.57 for SLBs and BSLBs, respectively. In other words, GP in DPPC:Chol GUVs is 0.65 GP units higher compared to POPC GUVs; however, GP in DPPC:Chol SLBs is only 0.44 units higher compared to POPC SLBs (Fig. 5E). These data highlight a tight and near-linear relationship between ordering and diffusion in GUVs but not in supported membranes which exhibit decreased sensitivity to compositional changes and deviation from the tight relationship between ordering and diffusion.

## Discussion

There is growing interest in reconstituting membrane associated processes *in vitro* using model membrane systems given their proven potential to refine mechanisms and glean new hypotheses^6^. However, few studies have ventured a comprehensive comparison of their biophysical properties as a function of the inherent parameters (a support presence, its material, model geometry etc.)^18, 25, 28, 40-43^. Indeed, a few studies have highlighted this concern by demonstrating altered protein functionality in different membrane models^44, 45^. We believe, our comprehensive comparison of lipid diffusion and ordering between GUVs, planar SLBs, spherical BSLBs and GPMVs would be a guideline for future studies that can use similar approaches for different biophysical properties.

Employing super-resolution STED-FCS, we confirmed the presence of nanoscale hindrances that seemingly influence the relationship between lipid ordering and diffusion in supported models, contrasting the linear relationship we observed in free-standing GUVs. Lipid ordering was comparable between GUVs, SLBs, and BSLBs of equal composition – at least at the microscopic scale, suggesting a more direct influence of support on diffusion. Indeed, the drag imparted by a lipid support is widely appreciated^18, 27^. However, our application of STED-FCS adds a spatial-temporal framework for these interactions and suggests repeated, but brief interactions between support and constituting lipids. This importantly highlights the necessity of applying super-resolution techniques to better elucidate the bilayer structures and local physicochemical properties. Nanoscale surface perturbations due to the support material have also been previously appreciated to influence diffusion^27^. Therefore, it is possible that nanoscale changes in surface topography of supported models effectively ‘slow’ lateral diffusion. Our observed discrepancy with BSLBs could reflect a change of support material, in which application of other methodologies such as atomic force microscopy could prove most useful.

The influence that bilayer ordering poses over lateral mobility can essentially be explained by increased Van der waals efficiency between lipids^29, 35, 46, 47^. However, absent from previous diffusion studies is direct comparison with lipid ordering we employ here. When combined with the capacity of the GUVs to retain unhindered lipid dynamics, we were able to demonstrate a direct correlation between lipid ordering and diffusion. Indeed, this is an attractive property of GUVs; due to their lack of support, GUV diffusion is considered ‘free’ or, more importantly, predictable. The potential of model membranes is impressive given their inherent skeletal framework, and indeed current applications prove this^6, 7, 48-50^. Yet, our results highlight an unpredictability in model behaviour that can accompany the incorporation of design motifs, in this case the presence of a support, and should not be overlooked in sensitive analysis.

## Acknowledgements

We thank the Wolfson Imaging Centre Oxford and the Micron Advanced Bioimaging Unit (Wellcome Trust Strategic Award 091911) for providing microscope facility and financial support. We acknowledge funding by the Wolfson Foundation, the Medical Research Council (MRC, grant number MC_UU_12010/unit programmes G0902418 and MC_UU_12025), MRC/BBSRC/EPSRC (grant number MR/K01577X/1), the Wellcome Trust (grant ref 104924/14/Z/14), and Wellcome Institutional Strategic Support Fund (ISSF). ES is funded by the Newton-Katip Celebi Institutional Links grant (352333122) and SciLifeLab fellowship. DB is grateful for the support of Integrated Immunology MSc program of University of Oxford.

## TOC Figure

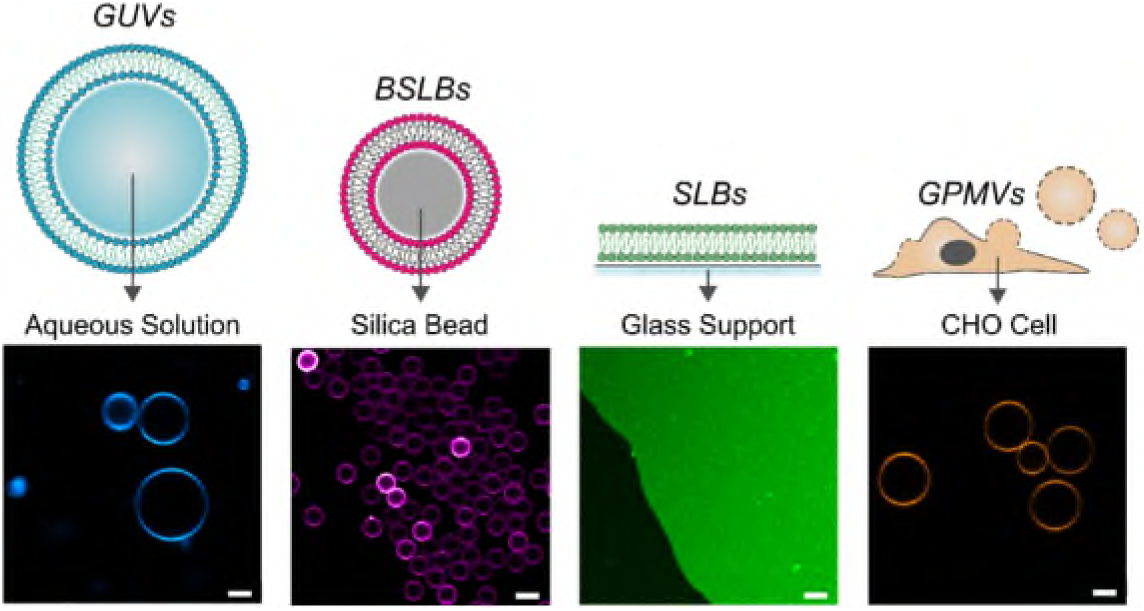

**TOC Figure Legend:** Comparing biophysical properties of model membranes

